# Integrated Approach to Investigate Magnetite Cycle in Marine Methanic Sediments

**DOI:** 10.1101/2025.03.30.646174

**Authors:** Adi Rivlin, Alice Bosco, Valeria Boyko, Ron Shaar, Yakar Zemach, Barak Herut, Susann Henkel, Keren Weiss-Sarusi, Orit Sivan

## Abstract

Magnetite (Fe_3_O_4_), a ubiquitous sedimentary iron mineral, is crucial for paleomagnetic records preservation. However, reactive ferric iron minerals, including magnetite, can undergo reduction in aquatic sediments above and within the sulfidic zone and at the Sulfate-Methane Transition Zone (SMTZ), resulting in the production of dissolved ferrous iron. Partial reoxidation of the reduced iron at the oxic-anoxic interface can lead to authigenic magnetite precipitation. Yet, magnetite persistence and behavior in deeper methanic sediments have remained poorly understood. Here we explore magnetite dynamics in different sub-methanic zones (deeper, middle and upper) of Mediterranean continental shelf sediments, including the potential for its authigenic precipitation. Sequential extractions revealed increasing magnetite concentrations accompanied by low-temperature magnetization (Verwey transition). First-order reversal curve (FORC) analyses supported nanoscale authigenic magnetite presence. The results highlight a net increase in single-domain magnetite in the middle methanic zone with declines in the upper and deeper zones. Microbial analyses pointed to iron reduction throughout the methanic zone, alongside potential precipitation of magnetite and a decline in methanogenesis functional genes. Sediment incubations with spiked ^57^Fe-ferrihydrite showed gross precipitation of isotopically enriched ^57^Fe-magnetite in the upper methanic zone. Our combined findings distinguish between gross and net magnetite precipitation, suggesting authigenic magnetite formation within the methanic zone, with reshaping and smoothing of the original magnetic signal. They also emphasize the limitations of relying on a single-method approach to unravel such complex processes. We propose that the methanic zone plays a critical role in the early diagenesis of magnetic minerals, driven by dynamic cycles of magnetite dissolution and authigenic precipitation.

## 1. Introduction

Microbial respiration of organic matter in marine sediments follows a cascade of electron acceptor reduction with decreasing free energy yield (Froelich et al., 1979; Jørgensen, 2000). Anaerobic respiration usually begins just below the oxygen-limited water-sediment interface, with nitrate reduction (denitrification), followed by manganese and iron oxide reduction, sulfate (SO_4_ ^2-^) reduction, and ultimately methanogenesis as the terminal step (Jørgensen, 2000; Roberts, 2015). In marine sediments, sulfate reduction coupled with anaerobic oxidation of methane (AOM) in the sulfate-methane transition zone (SMTZ) can consume large portion of the sulfate and up to 90% of the greenhouse gas methane produced by methanogenesis (Knittel & Boetius, 2009; Hoehler et al., 1994).

Below the SMTZ, the marine methanic zone is associated with reactive iron minerals, hosting active iron reduction (Aromokeye et al., 2020; Sivan et al., 2007; Vigderovich et al., 2019). The iron cycle in this zone can be linked to several processes, including AOM coupled to iron reduction (e.g. Sivan et al., 2007; Egger et al., 2015; Egger et al., 2017; Aromokeye et al., 2020), organoclastic iron reduction by dissimilatory iron-reducing bacteria (DIRB) (as shown in other scenarios by Conrad, 1999; Roden and Wetzel, 2003); iron reduction by FeS or pyrite oxidation (Amiel et al., 2020); and the metabolic switch of methanogens to iron reduction (Bond et al., 2002; Sivan et al., 2016; Eliani-Russak et al., 2023). Yet, the mechanisms and dynamics of iron cycling in this environment remain unclear.

The prevalence of a methanic diagenetic iron cycle poses a significant challenge to the paleomagnetic-dating research field (Amiel et al., 2020; Lin et al., 2020; Zemach et al., 2024), which relies on continuous and ideally undistorted records over geological time scales with global distribution (Panovska et al., 2019). Sedimentary paleomagnetic records are based on depositional remanent magnetization (DRM), where detrital magnetic particles are statistically aligned with the Earth’s magnetic field (Johnson et al., 1948). However, these records are vulnerable to distortion by diagenetic processes, including detrital input and the formation of authigenic iron magnetic minerals, which are influenced by porewater chemistry (Roberts, 2015).

In this work, we focus on magnetite (Fe_3_O_4_) as an indicator for iron cycling in the methanic zone. Magnetite plays a dual role as an electron acceptor in microbial respiration and a product of microbial iron reduction (Lovley et al., 1987; Kostka & Nealson, 1995). As an electron acceptor, the well crystalline magnetite represents the least favorable iron oxide of the reactive iron oxides, as iron reduction is strongly dependent on the Fe(III) level of crystallization (Thamdrup, 2000). It can also be formed extracellularly through dissimilatory iron reduction or intracellularly in magnetotactic bacteria as magnetosomes (Frankel, 1991; Chang & Kirschvink, 1989; Gareev et al., 2021). The behavior of magnetite and the potential for authigenic magnetite precipitation within the methanic zone are investigated here through a combination of geochemical, microbial, isotopic and magnetic methods.

Despite its potential significance, the role of the methanic zone in magnetic mineral diagenesis remains unexplored. Previous studies showed precipitation of the reduced iron as siderite (FeCO_3_) or vivanite (Fe_3_(PO_4_)_2_), paramagnetic minerals that do not gain magnetic properties. Profiles from Mediterranean Sea continental shelf sediments suggest that magnetite precipitation is linked to "deep" iron reduction (Amiel et al., 2020). However, experimental validations are limited, and identifying authigenic magnetite *in situ* remains challenging due to the small size of single-domain grains and limitations of existing techniques (Mathuriya, 2016; Liu et al., 2012; Lin et al., 2020).

Southeastern Mediterranean shelf sediments were deposited during the Holocene. Some settings feature a distinct shallow SMTZ at a depth of few meters, allowing us to define sub-zones within the methanic zone below it in 6–8-meter cores. Sediments were collected for detailed chemical and magnetic analyses, including porewater analyses of sulfate, methane, and dissolved iron. Mud slurries from the sub-zones were also incubated under various laboratory conditions to explore magnetite precipitation and/or dissolution. Magnetite detection in the sediment core as well as in the incubation experiments was performed using a combination of sequential extraction techniques and magnetic measurements. As these methods offer varying sensitivity to magnetite, the combined data enabled investigation of iron cycling in the methanic zones and was further contextualized by microbial DNA analyses. It should be noted that detrital magnetite is typically titanium-bearing and larger in size, while authigenic magnetite often forms as nanoscale grains (<30 nm) with superparamagnetic or single-domain properties (Ludwig et al., 2013). These distinctions are critical for paleomagnetic records, as diagenetic magnetite alteration can distort DRM records (Roberts, 2015).

## 2. Material and methods

### 2.1 Sampling sites

The study was performed in the Israeli continental shelf of the Southeastern Mediterranean Sea. Sediment cores were taken from stations PC3 (32.9216°N, 34.9027°E; Zemach et al., 2024) and SG1 (32.5787°N, 34.5530°E; Sela-Adler et al., 2015; Vigderovich et al., 2019; Amiel et al.,2020), located 20 km offshore Haifa and Acre at water depths of 80-90 m. The sediment in this area is mainly muddy clay, containing relatively low total organic carbon (TOC) of ∼1% (Sela-Adler et al., 2015). The sedimentation rate for the PC3 core was calculated by Zemach et al. (2024) based on radiocarbon dating and was found to be 3.5-3.6 mm/yr covering time frame of 1.0±0.1 ky.

Six-to eight-meter-long sediment cores were taken during cruises on the R.V. Bat Galim with a Benthos 2175 piston corer at stations PC3 in 2019 and 2020 and SG1 in 2017 and 2021. Cores for geochemical profiles were sampled onboard in minutes after core retrieval. Porewater was extracted by rhizons in a ∼20 cm interval for sulfate, nutrients, dissolved inorganic carbon (DIC), its stable carbon isotope composition (δ^13^C_DIC_) and Fe^2+^ concentrations. Porewater samples for PC3 sections from June 2020 between 450-490 cm depth and SG1 between 330-500 cm depth were discarded due to contamination with sea water. Sediments were also sampled for CH_4_ and δ^13^C_CH4_ within minutes after the core retrieval. Additional sediment samples were collected for incubation experiments and subsamples were frozen for microbial characterization and iron oxides extraction and magnetic measurements. Sediments from PC3 were collected for mineral magnetic analysis every 20-40 cm. The samples were freeze-dried and packed in non-magnetic gelatin capsules.

### 2.2 Isotope probing incubation experiment setup

In the lab, within hours after sampling, slurry incubations were prepared with sediments from four subzones of the methanic PC3-2020 core interval: (A) upper methanic zone (240-255 cm core depth), (B) and (C) middle methanic zones (300-320 and 370-390 cm), and (D) deep methanic zone (520-540 cm). Approximately 7 grams of sediment were inserted into 25 ml incubation glass bottles for a 1:1 slurry with synthetic anoxic seawater not containing sulfate (Table 1). They were flushed three times with N_2_ to ensure anaerobic conditions, and 3 different treatments were done to test iron-methane relation (Table 1). The first treatment (T1) was spiked with ^13^C-labeled CH_4_ (^13^C-CH_4_) to monitor the transformation of ^13^C-CH_4_ into ^13^C-DIC as an indication for its oxidation and examine Fe-AOM existence. To the second treatment (T2) we added ^57^Fe-spiked ferrihydrite to trace its reduction to Fe^2+^ and potential subsequent precipitation as ^57^Fe-contaoning magnetite. Synthetization of ^57^Fe-ferrihydrite was performed using ^57^Fe powder and the protocol of Cornell & Schwertmann (2003). The third treatment (T3) involved the addition of ^13^C-CH_4_ and un-spiked magnetite to evaluate the reduction potential of magnetite in the sediments coupled to methane oxidation. For T2 and T3 treatments, an autoclaved killed control was included to assess the effects of abiotic processes (T4, T5).

**Table 1:**
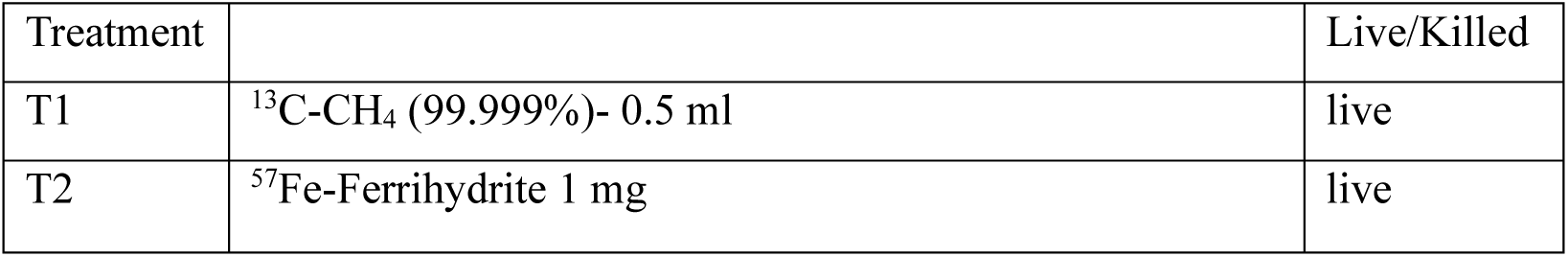
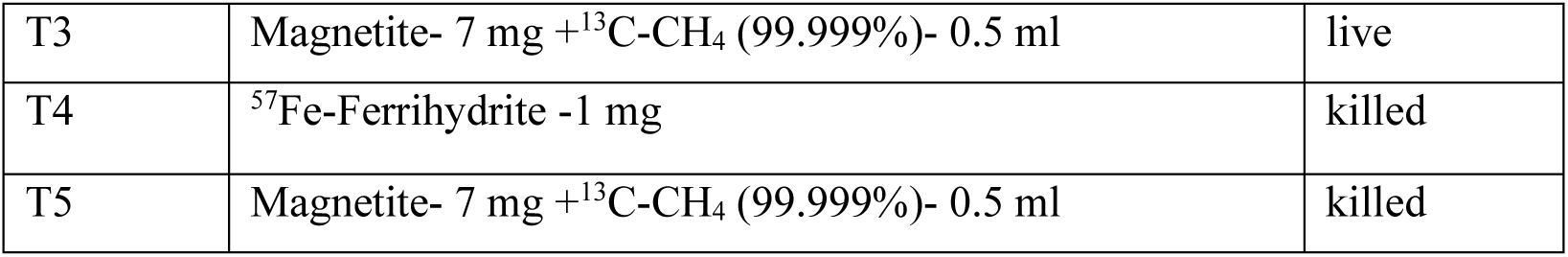
Details of isotopes probing incubation experiment from the Mediterranean Sea, station PC3.

Samples for dissolved Fe^2+^, DIC, δ^13^C_DIC_ and CH_4_ were taken measured at 5 time points: the start of the experiment (t_0_), after 7 days (t_1_), 31 days (t_2_), 96 days (t_3_) and at the end of the experiment after 130 days (t_4_). The sedimentary magnetite content, the magnetite isotopic composition, and magnetic parameters FORC and Verwey transition were determined for t_0_, t_2_ and t_4_. Samples for solutes were taken from the solute of the slurry through the rubber stoper with a needle and a 22 µm filter after vigorous shake and for CH_4_ from the head space. Afterwards, the incubation’s slurry was freeze-dried and processing of the sediment (sequential iron extraction, isotopic measurements, FORC and Verwey transition) started. Unfortunately, due to the large volume of sediments required for this type of experimental set up there was limited number of incubations, without replicates.

### 2.3 Analytical methods

#### 2.3.1 Porewater chemistry

For the measurements of methane concentration and its stable carbon isotopes (δ^13^C_CH4_) in the sediment depth-profiles, about 3 ml of sediment were collected and inserted immediately into 12 ml glass bottles filled with 5 ml of 1.5 mol L^-1^ NaOH and flushed with N_2_. The concentration of methane in the headspace of the bottles was measured with a focus gas chromatograph GC equipped with flame ionization detector (FID) at a precision of 2 μmol L^-1^. Sulfur concentrations were measured from the extracted porewater with Inductively Coupled Plasma Optical Emission spectroscopy ICP-OES. Sulfate concentration was calculated as the only contributing component to the sulfur concentration, as sulfide showed nano-molar concentration along the core and sulfite and thiosulfate showed micro-molar concentrations. For DIC concentration and δ^13^C_(DIC)_, porewater was added to glass tubes with phosphoric acid and measured with isotope-ratio mass-spectrometer IRMS equipped with a PreCon interface after oxidation to CO_2_. Isotopic measurements of DIC and methane are expressed in δ^13^C(‰)=10^3^((R_sample_ /R_standard_)-1), where R is ^13^C/^12^C and standards refer to the VPDB with an error of ±0.1‰ . Dissolved inorganic nutrients (NO_3_, NO_2_, NH_4_, PO_4_) were determined using a Seal auto-analyzer system with colorimetric detection (Ozer et al., 2022). The limits of detection, estimated as three times the standard deviation of 10 measurements of the blank (low nutrient aged seawater collected from off-shore surface at the Levantine Basin) for PO_4_, NO_2_+NO_3_ (NO_x_), NO_2_ and NH_4_, were 8, 80, 60, and 90 nM, respectively. Dissolved Fe^2+^ was measured with ferrozine (after Stookey 1970) by a spectrophotometer at 562 nm wavelength with detection limit of 1 µM .

#### 2.3.2 Iron oxide fractions

Sequential extraction of reactive iron oxides after Poulton and Canfield (2005) with citrate concentration based on Henkel et al. (2016). Sediments were freeze-dried and approximately 0.5 g of dry sediment reacted with 10 ml of a sodium acetate solution (1 M, pH 4.5) for 48 h in a shaker water bath at 50°C to dissolve the carbonates. The sample was then centrifuged, and the solution was pipetted into a separate vial for quantification. We then added 10 ml 1 M hydroxylamine-HCl in 25% [v/v] acetic acid to the residue for 48 h to dissolve FeO(OH) (ferrihydrite and lepidocrocite). More crystalline phases such as goethite, hematite and akageneite were extracted by reaction with sodium dithionite solution (50 g L^-1^) buffered to pH 4.8 with 0.35 M acetic acid or 0.2 M sodium citrate for 2 h. After collecting this fraction, the remaining sample was reacted for 6 h with ammonium oxalate (0.2 M, pH 3.2) to extract magnetite. All extractants were transferred into a falcon tube containing ferrozine solution and then measured spectrophotometrically.

Sulfide fractions were extracted from the wet sediments in a two-step distillation process after Canfield et al. (1986) using boiling HCl for monosulfides (FeS, ‘acid volatile sulfides’, greigite and mackinawite-like phases) followed by 1M chromium chloride (CrCl_2_) and concentrated HCl addition for Fe sulfides (FeS_2_, usually pyrite and pyrrhotite). The hydrogen sulfide released during the acid distillation process was trapped in 3% zinc-acetate. Sulfide concentrations were then measured colorimetrically after Cline (1969).

#### 2.3.3 Microbial analyses

Quantitative PCR and 16S rRNA gene V4 amplicon pyrosequencing DNA were extracted from the 2017 sediment core using a Power Soil DNA Kit (MoBio Laboratories, Inc., Carlsbad, CA, USA) following the manufacturer’s instructions. Copy numbers of selected genes were estimated with qPCR as described previously (Niu et al., 2017) using specific primers: Uni519f/Arc908R and bac341f/519r for archaeal and bacterial 16S rRNA genes, respectively, and mlas/mcrA-rev for the mcrA gene, which encodes the - subunit of methyl-coenzyme M reductase. The amplification efficiency was 94.5%, 106.3% and 92.4% for the archaeal 16S rRNA, bacterial 16S rRNA and the mcrA gene, respectively (the respective R2 of the standard curve was 0.998, 0.998 and 0.995).

#### 2.3.4 Iron isotope analyses of experimental incubations

The δ^57^Fe of the magnetite in the experiment were taken only as indication for the potential of gross production of magnetite due to the shortage in sampled sediment that did not enable incubations duplicates. Data is expressed in δ^57^Fe(‰)=10^3^((R_sample_ /R_standard_)-1), where R is the ratio of ^57^Fe/^54^Fe. Iron isotopic results represent the δ^57^Fe of the magnetite in the incubation, meaning only the oxalate fraction from the sequential iron extraction.

After sequential extraction, the oxalate extrat was processed for isotope analysis according to Henkel et al. (2016): The solution was oxidized with aqua regia (prepared using destilled HCl and HNO_3_) and H_2_O_2_ (suprapure), then evaporated and heated for 24 h at 140 °C. During heating, oxalate crystals condensated at the rim of the beakers were flushed back with ultra-pure water. Residues were re-dissolved in aqua regia and H_2_O_2_. After boiling for 2 h at 120 °C and evaporation, residues were re-dissolved in HNO_3_ and iron precipitation was performed by addition of H_2_O_2_ and NH_4_OH. The precipitate was redissolved in 6M HCl and a column chromatography was performed after Schoenberg and von Blanckenburg (2005): We used the BioRad anion resin AG1 X4 (200-400 mesh, chloride form). After separation, the Fe fraction was evaporated again, dissolved in 0.3 M HNO_3_, the concentration was measured by Thermo Scientific iCAP 7000 ICP-OES and matched to a concentration of 0.5 ppm Fe for for MC-ICPMS measurement (Teutsch et al., 2005).

Iron isotope measurements were performed on a Neptune multi-collector plasma source mass spectrometer (MC-ICPMS) instrument at MARUM. The abundant iron isotope ^56^Fe was measured together with ^57^Fe (which was added to the incubations) and ^54^Fe which the values are normalized to. The ^53^Cr isotope was simultaneously measured to monitor interferences of ^54^Cr on ^54^Fe, and the data corrected accordingly. Standard is referred to IRMM-014. An in-house standard was also used for monitoring accuracy (Henkel et al., 2016).

#### 2.3.5 Magnetic measurements

A detailed explanation of the experimental protocols and data analysis procedures for magnetic measurements is provided in Zemach et al. (2024), which also includes a comprehensive paleomagnetic analysis of core PC3. In this study, we briefly summarize the fundamental principles of the magnetic methods and their use as a proxy for estimating magnetite concentration.

In first order reversal curves of FORC analysis (Pike et al. 2001; Roberts et al., 2014), a sample is initially exposed to a strong saturation field. The magnetic field is then decreased to a specific reversal field (Br), after which the magnetization (M) is measured as the field (B) is ramped back up toward saturation. This process is repeated 600 times, with each iteration slightly reducing the reversal field (Fig. S1a, supplementary material). The resulting collection of curves is used to construct a FORC diagram, a contour plot of the magnetization second Derivative 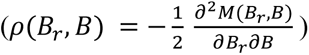 (Fig. S1b). The diagnostic behavior of sub-micrometer, non-interacting single-domain (SD) particles, known as the “central ridge,” appears in the FORC diagram as narrow, closed contours along the Bc axis of the FORC diagram. Egli (2013) and Egli et al. (2013) developed an algorithm to isolate the central ridge from the surrounding “background” signal, as shown in Figures S1c and S1d. We applied this isolation technique to calculate a magnetic proxy that quantifies the concentration of the smallest sub-micrometer ferrimagnetic particles relative to larger particles, by integrating the FORC diagram and normalizing it by the sample weight (Zemach et al., 2024). Assuming that detrital particles in the sediment are rarely sub-micrometer in scale, the central ridge magnetization serves as a proxy for authigenic magnetic minerals formed within the sediment, while the background magnetization reflects the concentration of larger, predominantly detrital magnetic particles.

Low-temperature measurements were conducted to identify the presence of stoichiometric magnetite by detecting the Verwey transition, characterized by a sharp decrease in magnetization between ∼85 K and ∼125 K (Jackson & Moskowitz, 2021; Walz, 2002). The specific temperature of the Verwey transition (Tv) is often used to infer the origin of the magnetite, with biogenic magnetite typically exhibiting Tv around 100K and inorganic magnetite around 120K (Chang et al., 2016; Jackson & Moskowitz, 2021). Measurements were performed using a Quantum Design Magnetic Properties Measurement System (MPMS3) in vibrating sample magnetometer (VSM) mode at the Institute for Rock Magnetism (IRM), University of Minnesota. The procedures (illustrated in Figure S2, supplementary material) included: warming a 2.5 T isothermal remanent magnetization (IRM) from 5K to 300K after cooling in zero-field (producing zero-field-cooled, ZFC, curves); warming of a 2.5 T saturation isothermal magnetization (SIRM) from 5K to 300K after cooling in zero-field (‘zero-field-cooled’, ZFC curves), warming of a 2.5 T SIRM from 5K to 300 K after cooling in 2.5 T field (‘field-cooled’, FC curves), cooling-heating cycle of room-temperature (RT) SIRM from 300K to 5K and back to 300K (SIRM cooling curve, SIRM heating curve). From these curves we calculated a proxy for the concentration of magnetite by taking the mass-normalized peak of the derivate between 100-120 K for each of the four above-mentioned curves.

## 3. Results

### 3.1 Dissolved iron and related porewater geochemical profiles

To identify the SMTZ and characterize iron dynamics within the methanic zone, geochemical porewater profiles were generated from two 6 m-long sediment cores collected at station PC3 during two campaigns in 2019 and 2020 (Fig. 2). We identified distinct differences in methane, dissolved Fe^2+^ and DIC concentrations in both campaigns, providing insights into the temporal variability of related geochemical processes in this sedimentary environment.

**Figure 1:**
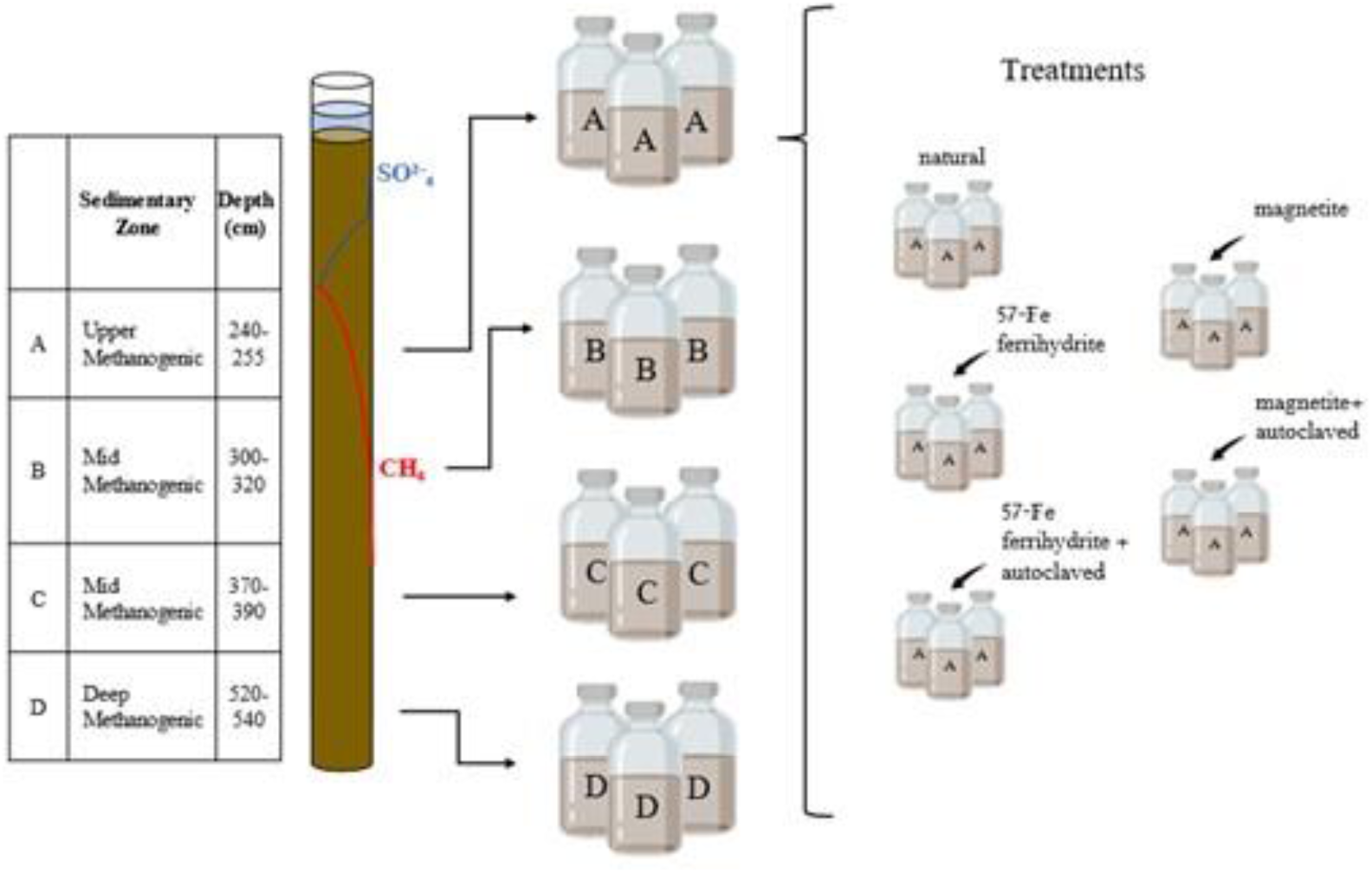
Incubation experiment set-up, table shows the depths of the methanic sediment which was taken for the experiment. For each depth the 5 different treatments were conducted and were measured for 5 time series of 0, 7, 31, 96 and 130 days.

**Figure 2:**
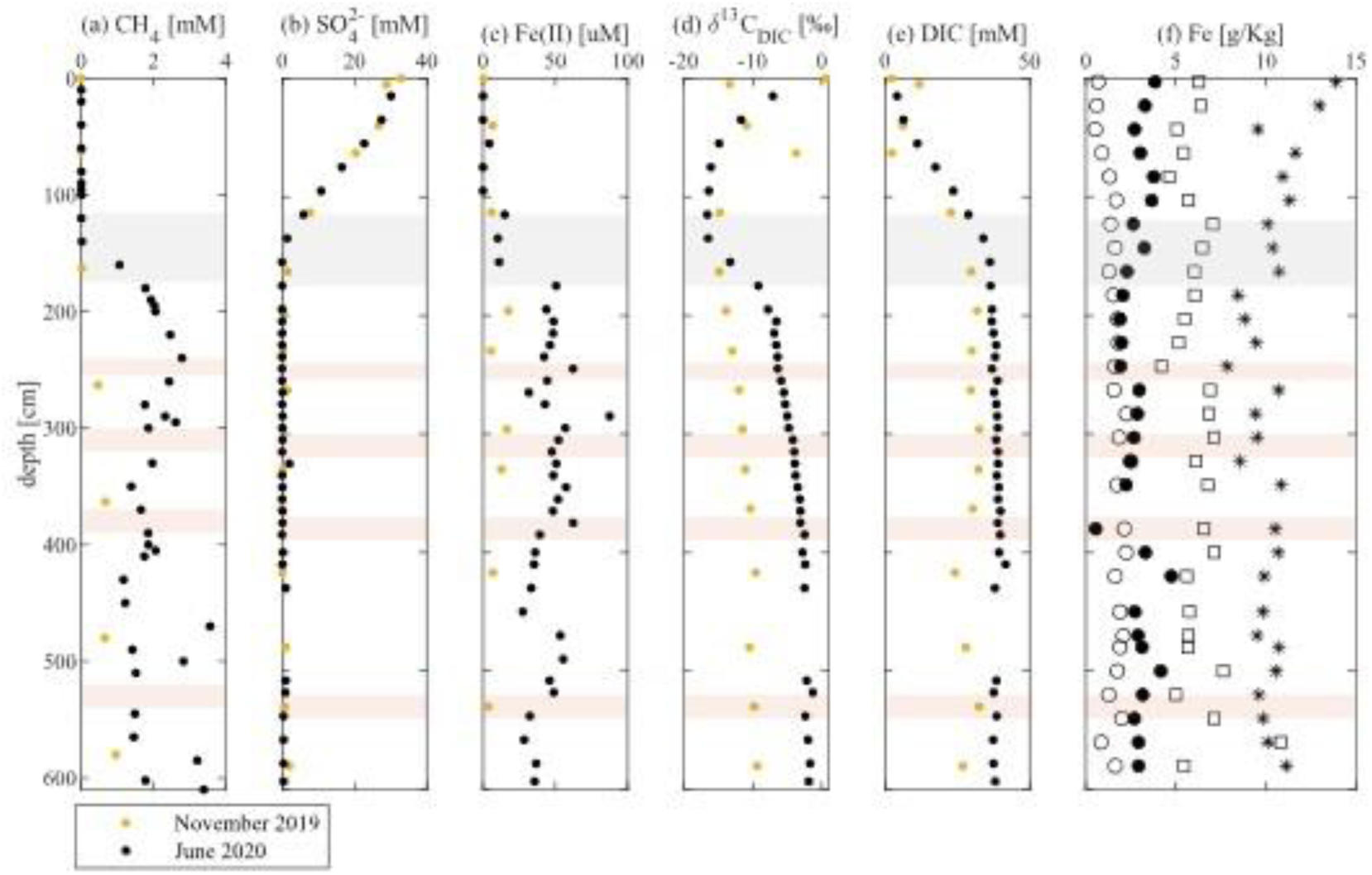
Porewater profiles from station PC3 in the Mediterranean Sea from two cruises in 2019 (yellow dots) and 2020 (black dots). Profiles show concentrations of (a) methane (CH_4_), (b) sulfate (SO ^2-^), (c) dissolved iron (Fe^2+^), (e) dissolved inorganic carbon (DIC), and (d) isotopic values of DIC (δ^13^C_DIC_ VPDB). In (f) Fe iron contents as determined by chemical extraction of Fe carbonates (empty circles), easily reducible Fe (empty squares), reducible Fe (asterisk) and magnetite (filled circles) are shown. Grey bars mark the SMTZ, and red bars (A to D) mark the four sections of the core from June 2020 which were taken for the incubation experiment.

During the 2019 campaign, sulfate concentrations decreased from 33 mM at the sediment-water interface to <1 mM at 195 cm. Methane concentrations increased to only 0.9 mM around 580 cm and Fe^2+^ concentrations in the methanic zone peaked at 17 µM. The DIC concentrations steadily increased, reaching maximum of 32 mM at the SMTZ with δ^13^C_DIC_ being light within the SMTZ (around −15‰), rising up to -9.4‰ around 600 cm. Sulfide concentrations were in nano-molar order of magnitude throughout the core, and are not shown because of near detection values.

In the 2020 campaign, SO_4_ ^2-^ concentrations decreased from approximately 30 mM at 15 cm depth to < 1 mM at 155 cm marking the SMTZ (∼120-170 cm). Methane started to accumulate at 160 cm reaching 2.7 mM at 240 cm depth and stabilizing around 1-2 mM down to 600 cm. Dissolved Fe^2+^ was minimal (<4 µM) in the upper sediments but increased significantly below the SMTZ, peaking at 87 µM at 285 cm depth and stabilizing around 50 µM throughout the methanic zone. DIC steadily increased from the sediment-water interface, reaching 38 mM in the SMTZ and remaining stable in the methanic zone. The δ^13^C_DIC_ were lighter within the SMTZ (around −16.5‰), rising up to −1.3‰ around 500 cm.

Reactive iron-bearing minerals from the PC3-2020 core (Fig. 2.f) showed that the content of Fe bound to carbonates ranged between 0.5 and 2.5 g kg^-1^ while Fe bound in easily reducible iron oxyhydroxides, minerals that are susceptible to microbial dissolution, ranged from 4.5 and 10 g kg^-1^. Iron bound in reducible iron oxyhydroxides contributed between 8 and 14 g kg^-1^ and Fe bound in magnetite was 3.8 g kg^-1^ at the core-top, decreased to 1.9 g kg^-1^ at 240 cm and showed notable increases below this depth. Peaks of magnetite-Fe at 420 cm (4.8 g kg^-1^) and 500 cm (4.1 g kg^-1^) exceeded the contents above the SMTZ, could suggest magnetite formation in the methanic zone.

Geochemical profiles from SG1 were obtained from 8 m long sediment cores collected as SG1-2021 and SG1-2017 (Fig. 3). As for PC3, the cores revealed distinct variations in sulfate, methane, dissolved iron, and other key chemical species. In SG1-2017, SO_4_ ^2-^ concentrations decreased from 31.5 mM at 10 cm depth, to <1 mM around 100 cm, defining the SMTZ from 100 to 150. Methane increased below 100 cm, reaching ∼1.5 mM at depths of 245 and 480 cm. Dissolved Fe^2+^ exhibited a peak above the SMTZ (∼85 µM), decreased to 5 µM at 125 cm and rose again to 65 µM at 245 cm. The DIC sharply increased from 2.8 mM at the core top to 27.3 mM at the SMTZ ∼50 cm, then steadily decreased to 6.4 mM at the core bottom (575 cm). The δ^13^C_DIC_ mirrored this trend, dropping from -2.9‰ at the surface to -25.8‰ at 170 cm, followed by an increase to −16.5‰ at the bottom core.

**Figure 3:**
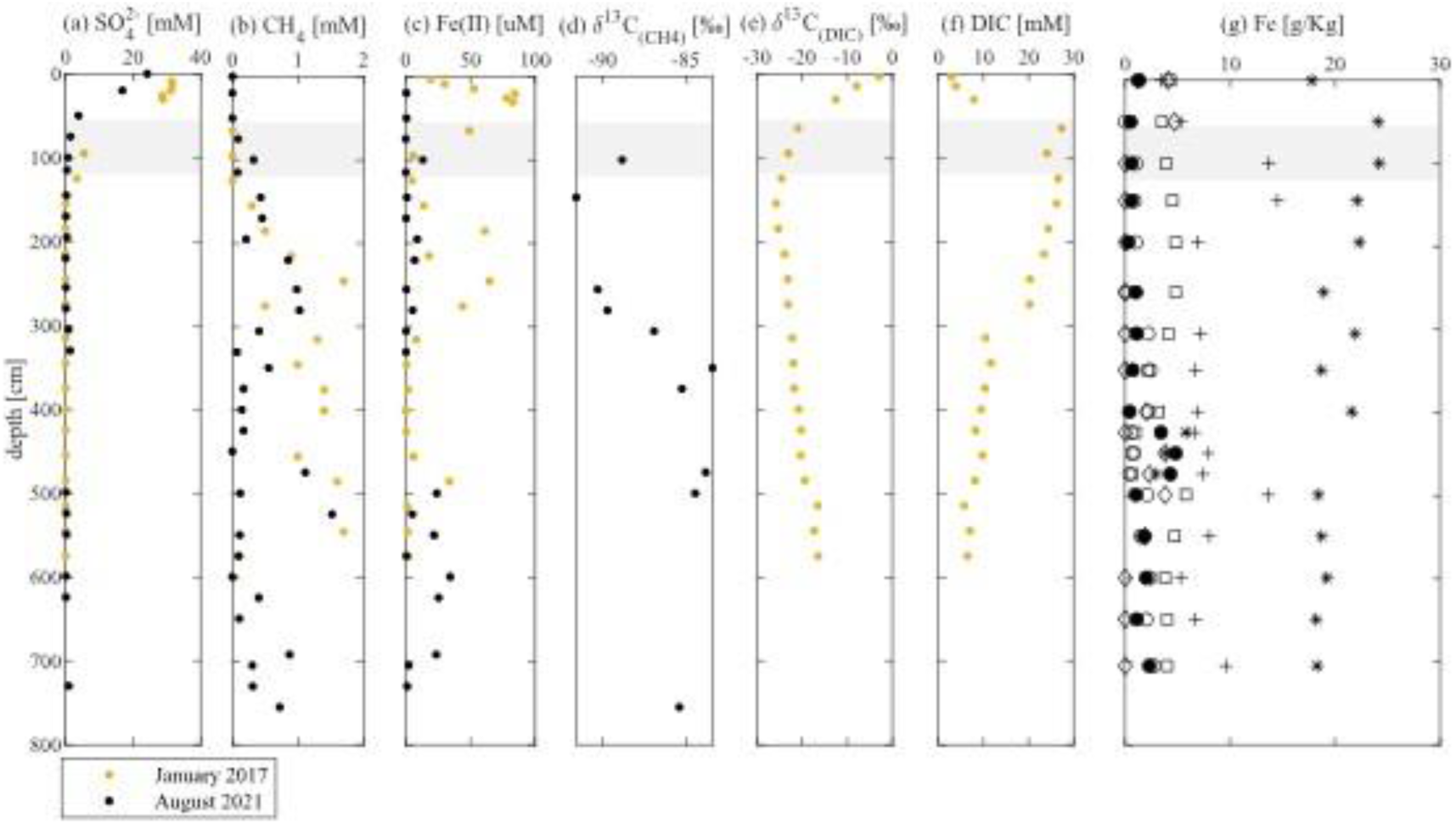
Porewater profiles from station SG1 in the Mediterranean Sea from cruises in January 2019 (after Vigderovich et al., 2019) and August 2021 (black dots). Profiles show (a) concentrations of sulfate (SO ^2-^), (b) methane (CH ), (c) dissolved iron (Fe^2+^) and (f) DIC. Isotopic values represented as δ^13^C (VPDB) of methane (d) and DIC (e). (g) Iron fractions profile: Fe carbonate (empty circles), easily reducible Fe (empty squares), reducible Fe (asterisk), crystalline oxide-magnetite (filled circles), acid volatile sulfur (empty diamonds) and chromium reducible sulfur (pluses). Grey bars mark the SMTZ.

In SG1-2021, SO_4_^2-^ concentration decreased from 24.1 mM at the sediment-water interface, to <1 mM around 150 cm, defines the SMTZ from 150 to 250 cm. Methane concentrations increased up to ∼1mM at 280 cm depth and ranged between 80 to 1114 µM within the methanic zone. The δ^13^C_CH4_ exhibited a minimum of -91.6 ‰ at 145 cm, and a maximum -83.8 ‰ at 474 cm. Dissolved Fe^2+^ concentrations increased below the SMTZ ranging from 0.1 to 13.1 µM between 20–330 cm, and further increasing to 34.3 µM below 500 cm. Concentration of NH_4_ ^+^ increased from 415 µM at 20 cm depth to over 1200 µM at the SMTZ and to a maximum of 2531 µM at 280 cm (Fig. S1). Concentration of PO_4_ ^3-^ increased from 1.4 µM at 20 cm to 4.94 µM in the SMTZ, in correspondence to NH_4_ ^+^ the maximum peak at 280 cm with 23 µM. Concentrations of NO_3_ ^-^ and NO_2_ ^-^ were unexpectedly detectable in the methanic zone with maximum values of 208 and 0.91 µM, respectively.

Iron fraction profiles (Fig. 3g) from the SG1-2021 core revealed contents of Fe bond in carbonates between 0.5 and 2.5 g kg^-1^, easily reducible iron oxyhydroxide between 0.7 and 5.8 g kg^-1^ and reducible iron oxyhydroxide between 2.9 and 24.0 g kg^-1^. The Fe derived from the magnetite concentrations were stable between 0-400 cm but showed a distinct peak of 4.8 g kg^-1^ between 400-500 cm, coinciding with decreases in other iron fractions except for iron-sulfur minerals. Below the peak contents were between 2.7 and 4.1 g kg^-1^. The AVS is relatively low with peaks above the SMTZ and in the methanic zone between 400-600 cm. Chromium-reducible sulfur, elemental sulfur and pyrite (FeS_2_), increased in the SMTZ up to 1.45 g kg^-1^ and then decreased in the upper methanic zone to 0.11 g kg^-1^ in 255 cm depth. The CRS below 300 cm depth stabilized at ∼0.6 g kg^-1^ with a distinct peak in 500 cm depth.

### 3.2 Magnetite cycle along PC3 sediment core

The magnetization depth plots derived from FORC analysis, reveal a growing population of non-interacting authigenic single-domain (SD) particles above the sulfate-methane transition zone (SMTZ) (Figure 4a-b). The Verwey transition proxies for magnetite, indicate that this authigenic phase is primarily composed of magnetite (Figure 4c-d). A notable decrease in magnetite concentration proxies is observed at the SMTZ, likely due to the reductive dissolution by hydrogen sulfide. However, the central ridge magnetization decreases less than the background magnetization. This may seem counterintuitive, as we would expect nanometer-scale particles (“central ridge”) to dissolve more rapidly than larger micrometer-scale particles (“background”). One possible explanation is the presence of SD ferrimagnetic greigite, which may form at or above the SMTZ. Another possibility is that SD magnetite continues to form within the upper methanogenic zone. Below the SMTZ, the magnetic data indicate detectable Verwey transitions due to magnetite throughout the entire methanic zone, consistent with the presence of a stable SD ferromagnetic population. The Verwey transition proxy also shows some fluctuations with depth, which may provide further support to the hypothesis of “deep” magnetite production.

**Figure 4:**
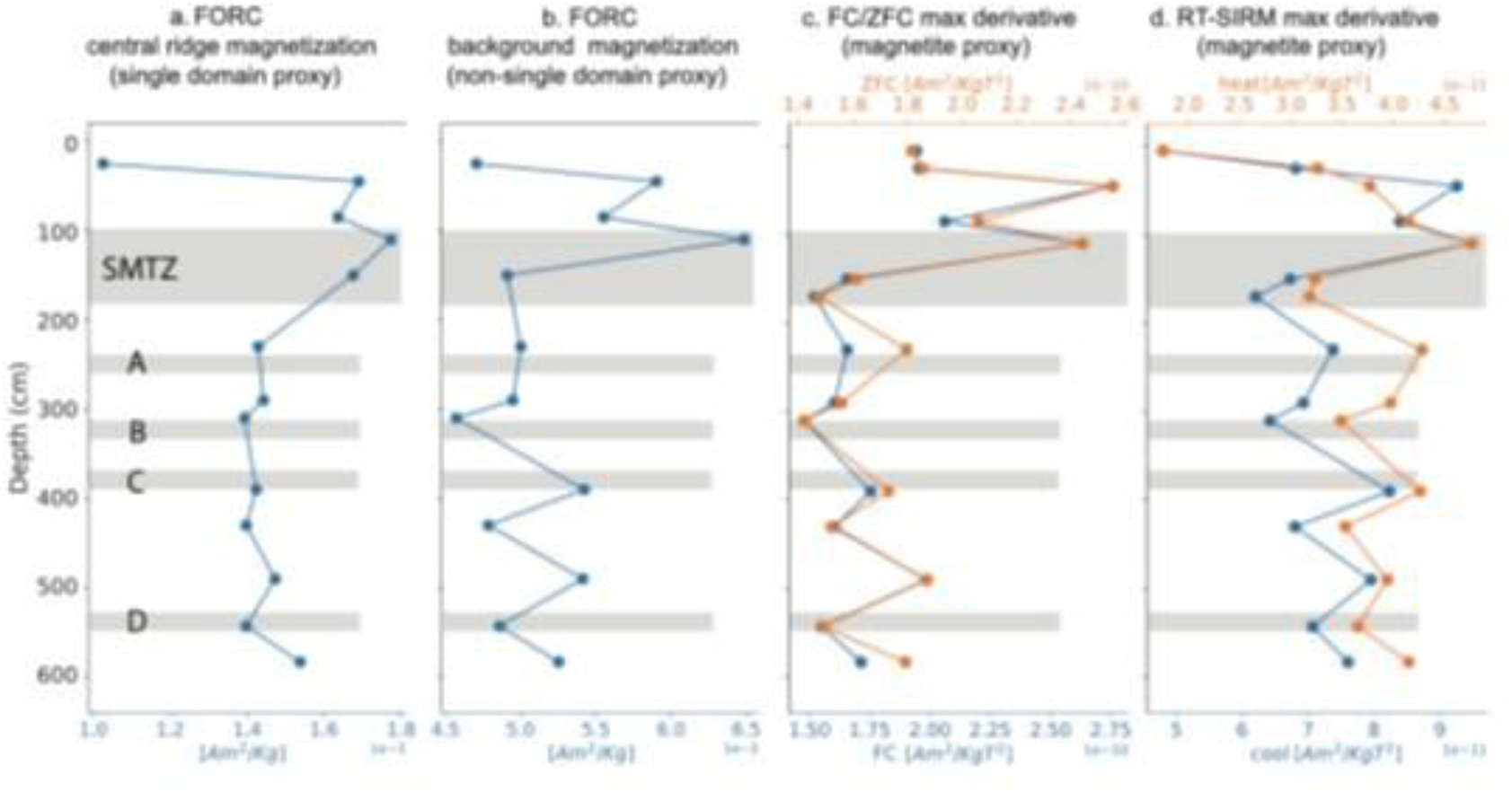
Depth variations of magnetic proxies in PC3. a-b) Mass-normalized magnetization calculated from the isolated central ridge and background of FORC diagrams, used as a proxy for the concentration of nanometer-scale single-domain ferromagnetic particles (a) and other particles (b). c-d) Magnetite concentration proxy calculated from the maximum derivative of the FC and ZFC cooled curves (c) and RT-SIRM cooling and heating curves (d), through the Verwey transition.

### 3.3 Microbial abundance along SG1 sediment core

Bacterial and archaeal abundance was measured along the sediment core from SG1-2017 (Vigderovich et al., 2019). Above the SMTZ bacterial community was mainly composed of a diversity of Chloroflexi (~34%, Dehalococcoidia, Ktedonobacteria, Thermomicrobia, Thermoflexia, and Anaerolineae) and Planctomycetes (~18%, mostly Phycisphaerae) groups (Figure S1). A decline of Proteobacteria (mainly Deltaproteobacteria from sulfate reducers families Desulfarculaceae, Desulfobacteraceae and Candidate Sva0485 clade) and increase of Deinoccocus-Thermus (from Deinococci class) were also observed at this depth interval. The Archaeal community was mainly composed of the Bathyarchaeota phylum, with abundance >70% throughout the entire core. The abundance of other archaea taxon, Thaumarchaeota and Euryarchaeota phylum, reached between 10-13% within the upper 100 cm.

At the SMTZ, Chloroflexi (42 to 27%), Deinoccocus-Thermus (21 to 9%), Planctomycetes (28 to 18%) and Proteobacteria (12 to 8%) were the most abundant bacteria phylum. The Nitrospirae community was reduced when compared to the sediments above the SMTZ, varying between 0.6 and 1.3% of the bacteria community. The bacteria phylum Elusimicrobia (known as magnetotactic bacteria (Gareev et al. 2022) reached 4% of the bacteria community at around 200 cm deep. The maximum relative abundances of Euryarchaeota from the anaerobic methanotroph ANME-1b group reached 10% of the archaeal community and 6.5% of the microbial community, followed by high distribution of mcrA gene compared to the other depths (> 30000 gene copies g-1, Fig. 5). This gene is highly positively correlated with FeS (Vigderovich et al., 2019), indicating AOM coupled to sulfate reduction (S-AOM), providing HS^-^ for iron sulfide precipitation.

**Figure 5:**
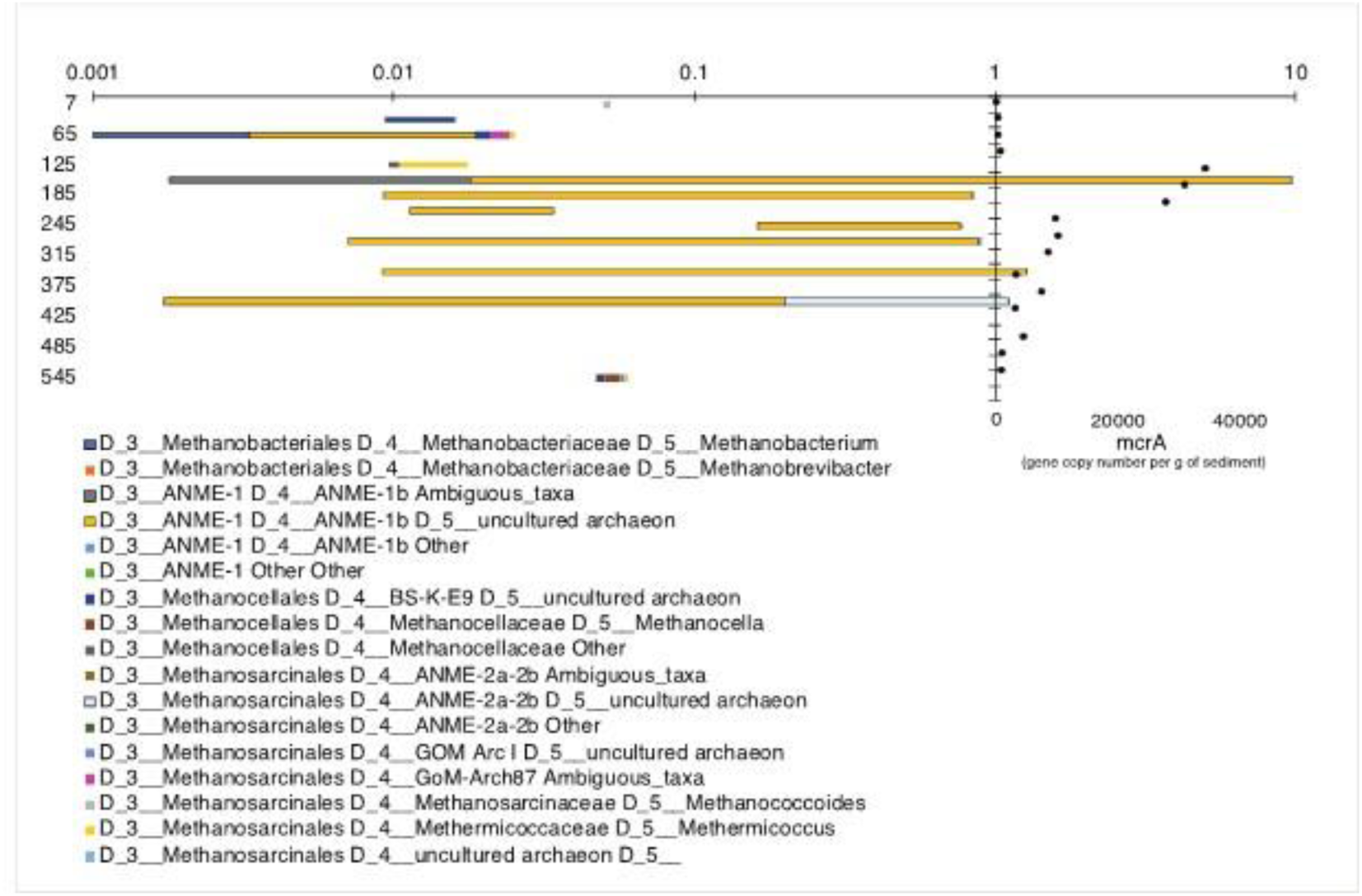
Relative distribution of methane-related archaea in the archaea community and gene copy number of *mcrA* per g of sediment by depth along the sedimentary profile from 2017.

Below the SMTZ Chloroflexi and Deinoccocus-Thermus abundance were smaller compared to the upper sediments (30 to 16% and 14 to 2%, respectively), while Planctomycetes and Proteobacteria abundances comparatively increased (32 to 18% and 15 to 11%, respectively). In the methanic sediments, increasing lots of Acidobacteria, Firmicutes, Defferribacteres and Spirochaetae were also observed. The archaeal community was dominated by Bathyarchaeota phylum, which has a relative abundance between 97 and 99%.

In the deep sediments, below 120 cm, ANME 2a-2b represents less than 2% of the archaeal community and 0.6% of the microbial community with low mcrA gene copies. This low microbial community contribution and of ANMEs that have not yet proven to participate in Fe-AOM hint that other processes would be involved in the reactivation of iron oxides reduction in the deep-sea sediments. At lower taxonomic levels and considering archaea methanotrophs from ANME in abundance were no higher than 0.6% of the microbial community at deep sediments, the most important dissimilatory iron reducers from Deferribacteres and Deinoccoccus-Thermus phylum show much lower abundance, respectively, at the methanic sediments.

### 3.4 Isotope probing incubation experiments

The DIC concentration and δ^13^C_DIC_ did not show a significant difference between treatment T1 with added ^13^C-CH_4_ and the magnetite added together with ^13^C-CH_4_ treatment (T3) at all depths (Fig. 6.2.a-d). Furthermore, δ^13^C_DIC_ of treatment T3 did not become heavier than δ^13^C_DIC_ of T1, suggesting no AOM (including Fe-AOM) occurred (Fig. 6.3.a-d). In sub-zone A, treatment T3 showed an increase in dissolved Fe^2+^ after 130 days to a maximum concentration of 94 µM, comparing to T1 and the killed treatment (T5) which accumulated only up to 73 and 65 µM (Fig. 6.1.a). When magnetite was added to the bottle it showed elevated iron reduction. Furthermore, methane production was inhibited in treatment T3 with the added magnetite (Fig. 6.4.a). In sub-zone B, Fe^2+^ and CH_4_ concentrations were similar in T1 and T3 treatments (Fig. 6.1.b and 6.4.b). In sub-zone C, when compared to treatment T1, magnetite treatment (T3) showed elevated iron reduction, but with a minor difference ranging between 5-10 µM (Fig. 6.1.c), and similar increases in CH_4_ (Fig. 6.4.c). In sub-zone D, treatment T3 showed lower concentrations of Fe^2+^ than treatment T1 (Fig. 6.1.d), and a similar increase in CH_4_ (Fig. 6.4.d).

**Figure 6:**
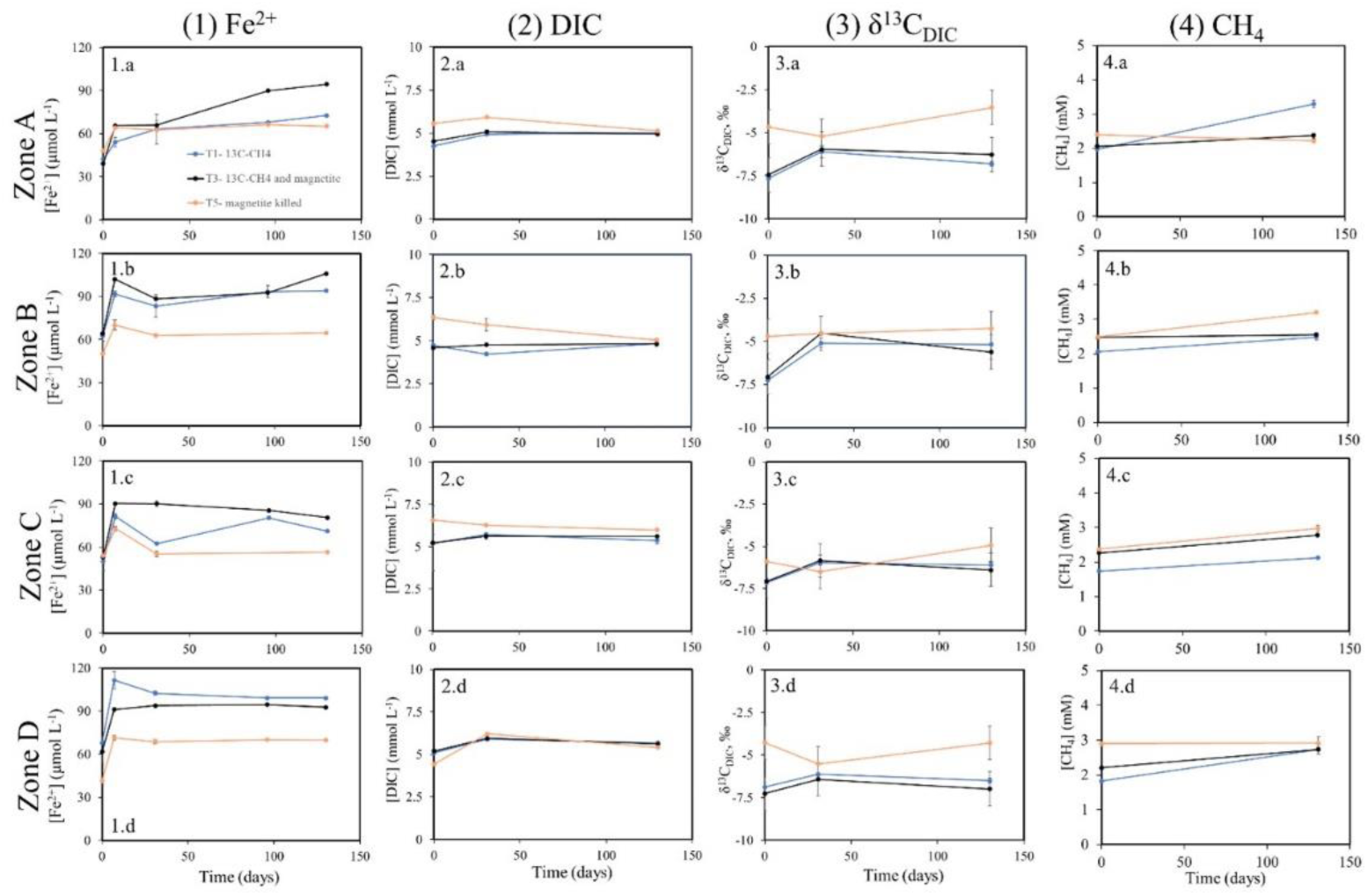
Dissolved species results of incubation experiment with methanic sediment from PC3 for depths A, B, C and D. The parameters were taken for treatment T1 with added ^13^C-CH_4_ (blue), treatment T3 with added magnetite and ^13^C-CH_4_ (black) and T5 magnetite and ^13^C-CH_4_ killed control (red): (1) dissolved Fe^2+^ concentration, (2) dissolved inorganic carbon (DIC) concentration, (3) the isotopic value δ^13^C_DIC_, and (4) CH_4_ concentration.

Changes in Fe-magnetite content in all experimental treatments over the time, using sequential iron extractions, are presented in Table S1 and show in general that with the sensitivity of the method of ±0.6 g kg^-1^, there was no significant change over time. The average content of magnetite-Fe in the beginning of the experiment in treatment T1 and in the ^57^Fe-ferrihydrite treatment (T2) was 2.8 g kg^-1^. The change (decrease or increase) after 130 days of experiment ranged between <0.1 and 0.4 g kg^-1^, and the average change in absolute value was 0.1 g kg^-1^, which equals the analytical error.

At t0, δ^57^Fe values were in probing method considered close to 0 (ranging from -0.24‰ to 28.01‰; Fig. 7.1.a-d) across all experimental setups, as expected. In the upper methanic zone (Fig. 7.1.a), the δ^57^Fe in treatment T2, with added ^57^Fe-ferrihydrite, increased from 18.90‰ at t2 to 25.31‰, and further to 313.74‰ at t4, indicating magnetite precipitation during the experiment. In contrast, no change in δ^57^Fe was observed in the middle methanic zone (Fig. 7.1.b-c) for treatment T2 from t0 to t4. In the deep methanic zone (Fig. 7.1.d), δ^57^Fe increased to 565.09‰ at t2 but returned to 25.49‰ at t4.

**Figure 7:**
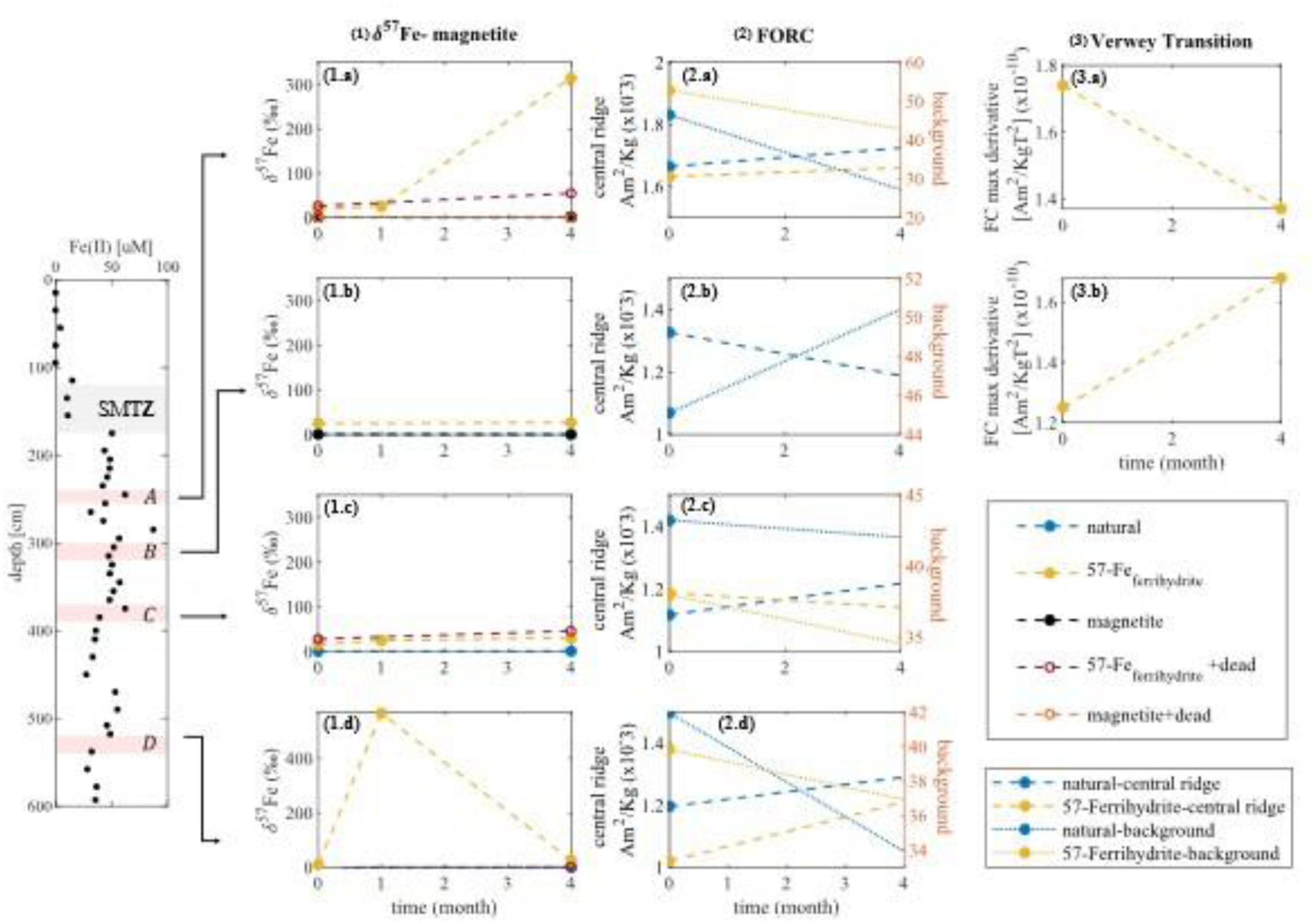
Incubation experiment using PC3 sediment core samples for time series of 0, 1 to 4 months. On the left Fe^2+^ profile for reference of sub-zones depth. (a) The δ^57^Fe values of magnetite fraction (oxalate extraction) for all treatments. Results for beginning and end of the experiment of the natural and ^57^Fe treatments with (b) FORC ’central ridge’ and ’background’, and (c) Verwey Transition, both normalized to sediment weight.

In treatments without ^57^Fe addition (T1, T3, T5), δ^57^Fe remained near 0 throughout the experiment, confirming no cross-contamination of the labeled isotope between treatments. The autoclaved ^57^Fe-ferrihydrite treatment (T4) showed an increased δ^57^Fe value of 28.33‰. However, the δ^57^Fe of the oxalate fraction was much larger (294.84‰) in the live treatment (T2), demonstrating that ferrihydrite-^57^Fe transitioned into the magnetite pool.

These results indicate gross magnetite formation in the upper methanic zone (Fig. 7.1.a) in the ^57^Fe-ferrihydrite treatment (T2). However, no isotopic changes were observed in the middle methanic zone (Fig. 7.1.b-c). In the deep methanic zone (Fig. 7.1.d), δ^57^Fe increased at t_2_ and decreased subsequently at t_4_ suggesting magnetite precipitation followed by its total consumption or dissolution. Alternatively, this pattern could result from analytical inconsistencies.

Both the ’central ridge’ and ’background,’ normalized to sediment weight, were measured at each depth for T1 with ^13^C-CH_4_ treatment (blue) and T2 ^57^Fe-ferrihydrite (yellow) treatments (Fig 7.2.a-d). In all sub-zones, except sub-zone C, in the ^57^Fe-ferrihydrite treatment T2 (Fig. 7.2.c), the ’background’ and ’central ridge’ displayed opposing trends.

In sub-zones A and D (Fig. 7.2.a, d), the concentration of SD ferromagnetic particles increased during the experiment, while other magnetic particles decreased in both treatments. In sub-zone B (Fig. 7.2.b), where only T1 was measured, SD ferromagnetic particles decreased, showing an opposing trend to the ’background’ phase. In sub-zone C (Fig. 7.2.c), the ’central ridge’ increased in T1 but decreased in T2, while the ’background’ phase decreased in both treatments.

Our results suggest that in sub-zones A, C, and D, new authigenic ferromagnetic minerals precipitated, while the ’background’ phase dissolved overall. In contrast, sub-zone B exhibited dissolution or growth of SD ferromagnetic minerals, accompanied by an overall increase in the ’background’ phase.

The Verwey Transition, normalized to sediment weight, was measured for sub-zones A and B in the ^57^Fe-ferrihydrite treatment (Fig. 7.3.a-b). This proxy for stoichiometric magnetite concentration showed detectable levels at the experiment’s start, with higher values in sub-zone A. Sub-zone A displayed a clear decrease over time, whereas sub-zone B showed an increase.

## 4. Discussion

In this study we investigated magnetite cycling in the methanic zone, challenging the long-held assumption that this zone is inactive regarding early diagenesis of magnetic minerals. Our results demonstrate significant magnetite dissolution and precipitation in the methanic zone, highlighting the dynamic nature of this environment and its implications for magnetic records and microbial survival in deep sediments. To address the inherent limitations of single-method approaches, we employed a multi-method strategy, integrating geochemical, isotopic, and magnetic analyses. Electron microscopy methods with high quality can identify authigenic magnetite but they are time consuming and not quantitative. Usually, the fraction of authigenic magnetite from the total magnetite is low, which makes it difficult to separate this pool, although the detrital magnetite is mostly found as titano-magnetite. There is no specific chemical procedure for distinguishing between the detrital and the authigenic magnetite. In addition, in sequential iron extractions, magnetite is partially extracted in the reducible Fe fraction. Similarly, in isotopic techniques, the difficulty in separating the authigenic magnetite from the rest of the magnetite in sediments limits the use of natural fractionation of stable iron isotopes. The magnetic tools also only provide a partial picture: The Verwey transition technique identifies the stochiometric magnetite, which is assumed to be authigenic, but is not sensitive for domain size. The FORC technique is sensitive for SD size, which is a proxy for authigenic minerals, but unspecific with regard to magnetite. Microbial diversity of bacteria and archaea were used only as supportive data that give indications for processes potentially occurring in the methanic zone. The isotope probing experiment provided evidence for **gross** authigenic formation, but the analytical challenges limit its use as well. The combined results are described below.

### 4.1 Evidence for reductive dissolution of magnetite in the methanic zone

The reductive dissolution of iron oxides including magnetite, a relatively low-reactive oxide, is traditionally not expected in the methanic zone, below the sulfidic zone. In addition, advanced thermodynamic and bioenergetic modeling indicates magnetite is the least favorable iron oxide for reduction in the methanic zone (Neumann Wallheimer, 2024). However, our multi-proxy results suggest magnetite reduction in the methanic zone of the continental shelf sediments in the Mediterranean Sea. In the incubation from sub-zone A, magnetite reduction was significant compared to the ^13^C-CH_4_ treatment and killed control, shown by Fe^2+^ accumulation in the incubation slurry (Fig. 6.2.a). This net dissolution was supported by FORC and Verwey Transition analyses, which revealed decreases in magnetite and a correlative decrease in the ’background’ signal (Fig. 7.b-c). Decreases in magnetite contents were also evident in deeper sub-zones (C and D), although less pronounced, and generally matched reductions observed in the natural control (Fig. 6.1.c-d). These findings are consistent with previous reports from the Mediterranean Sea (Vigderovich et al., 2019) and South China Sea (Liang et al., 2022), reinforcing the plausibility of magnetite reduction in the methanic zone. However, the exact drivers of magnetite reduction remain uncertain.

#### Ruled-Out Hypotheses

One proposed driver for magnetite reduction is methane, a potential electron donor abundantly produced in the upper methanic zone, as indicated by high *mcrA* gene copy numbers in the SG1 core (Fig. 5). Fe-AOM was evident in lake sediments and seep marine settings (Sivan et al., 2014). It has been also suggested as a potential mechanism in marine sediments (Sivan et al., 2007; Egger et al., 2017; Riedinger et al., 2014). However, our incubation experiments with ^13^C-labeled CH₄ showed no significant δ^13^C_DIC_ increase, effectively ruling out substantial Fe-AOM activity in the methanic zone (Fig. 6.c). This is further supported in this area by other sets of incubations (Yorshansky et al., 2022) and bioenergetic models (Neumann Wallheimer, 2024), and the absence of Fe-AOM-associated archaea, such as ANME-2a, in the sediments (Fig. 5). Low to insignificant Fe-AOM have been reported in the South China Sea (Liang et al., 2022) and the North Sea (Aromokeye et al., 2020).

Another hypothesis for magnetite reduction in the methanic zone involves DIRB, typically coupled with organic matter oxidation (Lovley & Phillips, 1988). However, the oligotrophic conditions of the Southeastern Mediterranean Sea suggest that most organic matter is oxidized above the SMTZ, leaving limited reactive organic compounds in the methanic zone. Hydrogen (H_2_) oxidation is another potential electron donor, given the measured H_2_ concentrations in SG1 (0.1–0.15 µM) and PC3 (2–6 µM) in Vigderovich et al. (2019). Nonetheless, incubation experiments with hematite and magnetite, with or without added H_2_, showed no significant difference in Fe^2+^ production (Vigderovich et al., 2019). This suggests that either endogenous H_2_ concentrations are sufficient for iron reduction or that H_2_ is not directly involved in this process.

#### Plausible Mechanisms for Magnetite Reduction

In the upper methanic zone, our data suggests that just below the SMTZ magnetite reductive dissolution may be driven by reactive sulfur species. These could include dissolved sulfide, FeS, or pyrite, which were not accounted for in thermodynamic models (Neumann Wallheimer, 2024). The near-complete pyrite consumption observed in the SG1-2017 core’s upper methanic zone supports a mechanism of pyrite oxidation coupled with magnetite reduction. This process coincides with low sulfate concentrations, likely due to pyrite oxidation or rapid sulfate consumption by methane or organic matter (Aplin & Macquaker, 1993; Bottrell et al., 2000).

Methanogens are also likely candidates for magnetite reduction. Studies with pure cultures of *Methanosarcina barkeri* have shown that methanogens can switch from methanogenesis to iron reduction, using ferrihydrite and magnetite as electron acceptors (Sivan et al., 2016; Eliani-Russak et al., 2023). Gene copy numbers of *mcrA* (Fig. 6) indicate that methanogens are concentrated in the upper methanic zone of SG1, which was also the most active zone for magnetite reduction, further supporting their involvement.

#### An Unusual Hypothesis: Feammox

A less conventional explanation for magnetite reduction is Feammox, the oxidation of ammonium (NH_4_^+^) coupled to iron reduction. In SG1, ammonium concentrations reached ∼2 mM in the methanic zone (Table S1), and surprisingly, nitrate (NO_3_^-^) was detected throughout the core, with a maximum of ∼200 µM at 170 cm in the upper methanic zone (Fig. S1). Given that Feammox typically oxidizes ammonium to nitrogen gas (N_2_) at the porewater pH, and Mnammox (ammonium oxidation coupled with manganese reduction) can produce nitrate, we propose that ammonium oxidation in this case involves both Feammox and Mnammox. Supporting this hypothesis, *Actinobacteria* from the *Acidimicrobiaceae* family, which includes the only known organisms capable of Feammox, were abundant in deep sediments, reaching up to 10% of the microbial community.

### 4.2 Authigenic magnetite precipitation in the methanic zone?

Authigenic magnetite is typically expected to precipitate above the SMTZ at the oxic-ferrous interface (Karlin et al., 1987) and to dissolve completely within the SMTZ due to its high reactivity with sulfide. However, studies have increasingly reported the reduction of iron oxides and the potential precipitation of authigenic magnetite in the methanic zone (Amiel et al., 2020; Lin et al., 2020; Zemach et al., 2024). Our results provide new evidence supporting magnetite cycling and potential authigenic magnetite formation in this previously underestimated zone.

In **sub-zone A** (upper methanic zone, 70 cm below the SMTZ), isotope probing revealed dynamic magnetite cycling. FORC data showed growth of single-domain (SD) ferrimagnetic minerals, likely not magnetite, as Verwey Transition indicated net dissolution of stoichiometric magnetite. However, δ^57^Fe values suggested magnetite precipitation during the experiment (Fig. 7.1-3.a). These findings indicate an active recycling process where magnetite undergoes both dissolution and precipitation. The simultaneous increase of other authigenic ferromagnetic minerals highlights iron oxide recycling as a key feature in this sub-zone.

In **sub-zone B** (main methanic zone), Verwey Transition detected an increase in stoichiometric magnetite, but FORC data showed dissolution of SD ferrimagnetic minerals. Isotopic results, however, revealed no evidence of new magnetite formation. This suggests that the observed increase in pure magnetite was due to crystal growth of pre-existing magnetite rather than authigenic formation of new grains.

**Sub-zones C and D** lacked comprehensive data due to analytical limitations. In sub-zone C (deeper in the main methanic zone), FORC analysis indicated a net increase in nano-sized ferrimagnetic minerals in the natural treatment, while the ^57^Fe-ferrihydrite treatment showed slight dissolution. However, δ^57^Fe data did not support magnetite formation during the experiment. In **sub-zone D** (deep methanic zone), FORC results suggested net formation or growth of nano-sized magnetic minerals, but isotopic issues prevented further interpretation.

Overall, the upper methanic zone (sub-zone A) appears to be magnetically active, with evidence pointing to authigenic magnetite formation and dynamic iron oxide cycling. Further work is needed to quantify the extent and significance of these processes, particularly in deeper sub-zones.

#### Microbial Involvement in Magnetite Cycling

The isotopic results strongly suggest microbial involvement in magnetite formation, with significantly higher δ^57^Fe-magnetite values observed in live treatments compared to autoclaved controls (Fig. 7.1.a). Microbes may contribute to magnetite formation through extracellular precipitation following iron-oxide reduction by DIRB, or via intracellular biomineralization by magnetotactic bacteria. The magnetotactic bacterium *Elusimicrobia* was identified in the sediments and was more abundant in the methanic zone, comprising up to 4% of the microbial community in deep sediments where magnetite is present. Methanogens, previously discussed as drivers of microbial iron reduction, may also play a role in extracellular magnetite formation (Lin et al., 2024).

#### Implications for Paleomagnetic Records

The discovery of authigenic magnetite in the methanic zone has significant implications for sedimentary paleomagnetic records. Newly formed magnetite can alter or "smooth" paleomagnetic signals, potentially complicating the interpretation of these records (Roberts and Weaver, 2005; Larrasoaña et al., 2007; Rowan et al., 2009; Roberts et al., 2018). These findings underscore the need to consider magnetite cycling in the methanic zone when interpreting magnetic records in marine sediments.

## 5. Conclusions

Our study emphasized the challenges in tracing the cycle of magnetite using a single method, and the limited information achieved. To overcome these difficulties, we used multidisciplinary approach. The contribution of each method is different and together provides a broader picture. Chemical iron extractions showed the variance of total magnetite (detrital and authigenic) concentration along the core, and that magnetite remained in the same order of magnitude in the methanic zone, with potential for reduction and precipitation. As mentioned, FORC ’central ridge’ analysis normalized to sediment weight is a proxy for authigenic minerals. When FORC results were compared to the Verwey transition results, in both sub-zones A and B, opposite trends were found. On the other hand, the trends of the ’background’ showed similar trends to Verwey transition. This implies that the contribution of magnetite to the increase of authigenic minerals concentration in the experiment is probably low. This challenges the assumption that magnetite is the main authigenic mineral contributing to the FORC ’central ridge’ signal observed in the methanic zone in PC3 profile (Fig 6.g). The similar trends of Verwey transition and FORC ’background’ but opposite to ’central ridge’ may indicate the growth of pure magnetite crystals, which can increase Verwey transition signature, rather precipitation of new authigenic magnetite. Combining and comparing FORC ’central ridge’ and ’background’ to Verwey transition analysis gave an indication for the nature of the authigenic minerals signal and the variation of pure magnetite signal.

Yet, these methods show net increase or decrease, which does not provide sufficient information on the gross reactions. Using probing method, addition of enriched isotopes, allowed for isotope ^57^Fe tracing and for the formation of magnetite in the slurry experiment. In this case even if there is net dissolution of magnetite, the enriched-isotope method may demonstrate the occurrence of this reaction. Nevertheless, this method presents numerous analytical challenges due to its complex procedures, which, in this study, resulted in a lack of repetition. This method showed precipitation of magnetite during the experiment in sub-zone A, unlike the other sub-zones. Microbial diversity of bacteria and archaea are supportive results, which give indications for microbial iron reduction by DIRB, methanogens or *Acidimicrobiaceae* by organic matter or NH_4_ ^+^, respectively.

The overall findings allow us to differentiate between gross and net magnetite precipitation, suggesting the potential for authigenic magnetite precipitation within the methanic zone. However, they also underscore the complexity of relying solely on a single method for such observations. We propose that the methanic zone is actively involved in the early diagenesis of magnetic minerals, characterized by dynamic processes of dissolution and precipitation of authigenic magnetite.

## Supporting information

Figure S1: Nutrients concentrations from SG1 core from 2021 cruise: Dissolved silica (Si(OH)4), phosphate (PO43-), nitrate (NO3-) nitrite (NO2-) and a

Table S1: concentrations of magnetite in the sediment during incubation experiment

## Notes

### Competing Interest Statement

The authors have declared no competing interest.

