## Supplementary figures and images for "Integrated Approach to Investigate Magnetite Cycle in Marine Methanic Sediments"

### Figure S1: Nutrients concentrations from SG1 core from 2021 cruise: Dissolved silica (Si(OH)4), phosphate (PO43-), nitrate (NO3-) nitrite (NO2-) and a

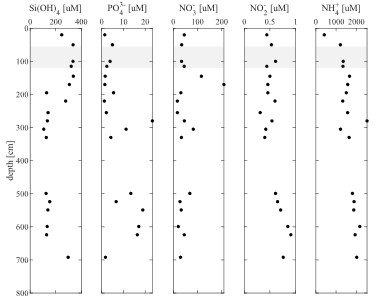
