## Supplementary material for "Integrated Approach to Investigate Magnetite Cycle in Marine Methanic Sediments": Table S1: concentrations of magnetite in the sediment during incubation experiment

| | | t0 | t1 | t2 |
|---|---|---|---|---|
| | | ±0.06 g/Kg Fe(magnetite) | | |
| A | natural | 2.84 | 2.85 | 2.83 |
| | $^{57}$Fe ferrihydrite | 2.83 | 2.92 | 2.92 |
| | $^{57}$Fe ferrihydrite+autoclave | 2.95 | 3.04 | 2.94 |
| B | natural | 2.82 | 3.03 | 2.94 |
| | $^{57}$Fe ferrihydrite | 2.89 | 3.10 | 3.29 |
| | $^{57}$Fe ferrihydrite+autoclave | 3.04 | 3.11 | 3.13 |
| C | natural | 2.88 | 2.63 | 2.76 |
| | $^{57}$Fe ferrihydrite | 2.91 | 2.77 | 2.83 |
| | $^{57}$Fe ferrihydrite+autoclave | 2.82 | 2.87 | 2.65 |
| D | natural | 2.76 | 2.61 | 2.75 |
| | $^{57}$Fe ferrihydrite | 2.82 | 2.86 | 2.72 |
| | $^{57}$Fe ferrihydrite+autoclave | 3.04 | - | 2.78 |
